# *Nasonia* segmentation is regulated by an ancestral insect segmentation regulatory network also present in flies

**DOI:** 10.1101/2021.03.23.436706

**Authors:** S E Taylor, P K Dearden

## Abstract

Insect segmentation is a well-studied and tractable system with which to investigate the genetic regulation of development. Though insects segment their germband using a variety of methods, modelling work implies that a single gene regulatory network can underpin the two main types of insect segmentation. This means limited genetic changes are required to explain significant differences in segmentation mode between different insects. Evidence for this idea is limited to *Drosophila melanogaster, Tribolium castaneum*, and the spider *Parasteatoda tepidariorum*, and the nature of the gene regulatory network (GRN) underlying this model has not been tested. Some insects, for example *Nasonia vitripennis* and *Apis mellifera* segment progressively, a pattern not examined in studies of this segmentation model, producing stripes at different times throughout the embryo, but not from a segment addition zone.

Here we aim to understand the GRNs patterning *Nasonia* using a simulation-based approach. We found that an existing model of *Drosophila* segmentation (Clark, 2017) can be used to recapitulate *Nasonia*’s progressive segmentation, if provided with altered inputs in the form of expression of the timer genes *Nν-caudal* and *Nν-odd paired*. We also predict limited topological changes to the pair rule network. Together this implies that very limited changes to the *Drosophila* network are required to simulate *Nasonia* segmentation, despite the differences in segmentation modes, implying that *Nasonia* use a very similar version of an ancestral GRN also used by *Drosophila*.

## 1 Introduction

Embryonic segmentation has been extensively studied, both genetically and embryologically, in a number of insects and represents an excellent system with which to understand the genetic regulation of development, and its evolution (see Akam, 1989; Davis and Patel, 2002; Peel and Akam, 2003; Clark et al., 2019; Chipman, 2020 for reviews). Insect segmentation is best understood in *Drosophila melanogaster,* where two protein gradients, of bicoid (bcd) and caudal (cad), establish the early anterior-posterior pattern. Proteins encoded by gap genes then subdivide the body axis into broad domains, and regulate expression of the pair rule genes. Expression of the pair rule genes marks the first periodic gene expression in the embryo, these genes regulate expression of the segment polarity genes, establishing the morphological segments (Jaeger, 2011; Clark et al., 2019).

Insect trunk segmentation is broadly classified into two types: simultaneous and sequential segmentation. Simultaneously segmenting insects produce each body segment at the same time, while sequentially segmenting insects produce new segments one after the other, from a posterior segment addition zone (Davis & Patel, 2002; Peel & Akam, 2003; Clark et al., 2019). This simultaneous/sequential classification is an over-simplification. *Nasonia vitripennis,* the subject of this study, and the honeybee *Apis mellifera,* express segment polarity and pair rule genes in an anterior to posterior progression (Bull, 1982; Fleig & Sander, 1988; Binner & Sander, 1997; Osborne & Dearden, 2005; Wilson & Dearden, 2012; Rosenberg et al., 2014). That is, new segments are specified one after the other, but are not produced from the posterior of the embryo. This type of segmentation, *progressive segmentation*, is an intermediate between simultaneous and sequential segmentation. *Nasonia* combine progressive segmentation in the anterior with sequential segmentation in the posterior, implying that progressive and sequential segmentation can be achieved in the same species and presumably using the same genetic background (Rosenberg et al., 2014).

Modelling and empirical work implies that simultaneous and sequential segmentation can be achieved using the same molecular process. The *Drosophila* pair rule GRN is best understood as two networks, early and late (Clark & Akam, 2016). The early network produces periodic, pair rule gene expression, while the late network converts this pair rule pattern into the segment polarity pattern (Clark, 2017). Changing the timing of network activation controls whether segmentation is simultaneous or sequential. If the timing of network activation is the same across the whole embryo, simultaneous segmentation occurs as each segment matures at the same time. If the networks are activated at different times throughout the embryo, the embryo segments sequentially (Clark, 2017; Clark & Peel, 2018).

Transitions between these networks are thought to be controlled by the timer genes *cad*, *Dichaete* / *sox21b*(*D*) and *odd-paired*(*opa*). *Cad* has a well-conserved role as a posterior determinant, and regulates pair rule gene expression in many species (Copf et al., 2004; Wilson et al., 2010; Kimelman & Martin, 2012; El-Sherif et al., 2014; Rosenberg et al., 2014; Schönauer et al., 2016; Zhu et al., 2017; Novikova et al., 2020). *D* is required for normal expression of some pair rule genes in *Drosophila* and for spider segmentation, and the expression of *D* orthologues in the segment addition zone is conserved throughout panarthropoda (Nambu & Nambu, 1996; Russell et al., 1996; Ma et al., 1998; Clark & Peel, 2018; Janssen et al., 2018; Paese et al., 2018; Baudouin-Gonzalez et al., 2020). In *Drosophila, opa* is required for late network activation and genome-wide regulatory changes (Clark & Akam, 2016; Koromila et al., 2019; Soluri et al., 2019). In other insects, *opa* is expressed in a band at the end of the segment addition zone, where a putative late network would be active, though *Oncopeltus* lack this pattern, implying this timer gene role may not be conserved across all insects (Green & Akam, 2013; Xiang et al., 2017; Janssen et al., 2011; Auman & Chipman, 2018). Together, this correlative evidence implies that the timer genes *cad*, *D*, and *opa* are plausibly involved in regulating segmentation across many arthropod species.

The timer gene proposal has important implications. Firstly, it provides a simple way to evolve phenotypic diversity: expressing the timer genes in different patterns causes different activation of the pair rule networks, and a switch between sequential and simultaneous segmentation (Clark, 2017; Clark & Peel, 2018; Clark et al., 2019). Secondly, it provides a biological example of a multifunctional GRN. The multifunctionality of GRNs modelled as ordinary differential equations is well-documented. Most dramatically, the AC/DC circuit (three genes in a circuit, each repressing the one after, and one pair of genes also repressing each other) is capable of both oscillations and multistable behaviour (Panovska-Griffiths et al., 2013; Perez-Carrasco et al., 2018; Verd et al., 2019). These different behaviours are central to explaining the evolvability of the gap gene system of flies (Verd et al., 2019). Also, gene circuits with the same topology are capable of different dynamical behaviours when the weights of interactions between genes are changed (Jiménez et al., 2017). The pair rule system provides another example of a GRN which is capable of two functions: sequential and simultaneous segmentation. Both of these types of segmentation rely on the same dynamic behaviour of the GRN: the ability of the early network to oscillate within a cell. The behaviour of the GRN is unchanged, but how it is used by the organism is different (Wouters, 2003).

The lack of correspondence between a GRN topology and its dynamical behaviour means that we cannot analyse GRNs based on their structure alone (DiFrisco & Jaeger, 2019). Instead we should dynamically model GRNs to understand them, as Clark, 2017 does for *Drosophila* (DiFrisco & Jaeger, 2019). Such models are challenging to construct from the ground up for non-model species, requiring impressive amounts of experimental data which are typically lacking (Peter & Davidson, 2016; Crombach et al., 2012; Wotton et al., 2015; Clark, 2017). Moreover, simply identifying which genes are involved in a given process, via functional data and analysis of gene expression patterns, does not give sufficient information into how these genes may interact, and how the network as whole behaves to produce patterning.

Here, we attempt an alternative approach. We use hybridization chain reaction (HCR) to describe the process of *Nasonia* segmentation in spatial and temporal detail (Choi et al., 2016; Choi et al., 2018). We then use an existing model of *Drosophila* pair rule patterning (Clark, 2017) to identify where this model can and cannot account for observed changes in *Nasonia* gene expression. We use this simulation-based approach to predict where changes in gene regulation are required to produce *Nasonia-like* patterning, and where the altered *Nasonia* inputs are sufficient to produce the observed changes. This computational model idealises the embryo as a one-dimensional row of cells, obeying boolean logic to determine how gene expression (protein and RNA expression and age) changes over time (Clark, 2017).

We find that surprisingly few changes to the *Drosophila* pair rule GRN are required to simulate *Nasonia-like* patterning, implying that there may be limited topological changes to this network throughout insect evolution.

## 2 Results

### 2.1 Dynamics of gene expression

To compare *Nasonia* and *Drosophila* gene expression during segmentation, we needed a comprehensive and precisely staged description of *Nasonia* segmentation. To enable robust temporal and spatial characterisation of gene expression, we compared expression of each pair rule gene to the expression of *Nv-eve* and *Nv-wg*, alongside *Nv-sim* (a marker of the ectoderm boundary, Buchta et al., 2013) for selected genes. This allowed us to stage each embryo by the number of *eve* and *wg* stripes, and robustly characterise the relative timings of changes in gene expression. Stages are named by the number of wg/eve stripes: ie “eve5” embryos have five *eve* stripes. “Stripes” mean regions of the embryo expressing the gene of interest.

Like Rosenberg et al., 2014, we identified three distinct regions of the *Nasonia* embryo. In the anterior (eve stripes 1-5), *Nasonia* undergo progressive segmentation. Pair rule stripes of *Nv-hairy*, *Nv-odd*, *Nv-E75A*, and *Nv-ftz* are expressed in an anterior to posterior progression. The expression of *Nv-runt* and *Nv-eve* stripes is simultaneous: the first three stripes appear at around the same time, with *Nv-runt* being expressed first (around the same time as *Nv-odd*). As in *Drosophila*, *Nv-slp* is not expressed until after other pair rule genes, at the eve4 stage. Unlike *Drosophila*, *Nv-prd* is not expressed until the eve8 stage, after frequency doubling (Keller et al., 2010). Cellularization does not commence in *Nasonia* until at least the eve8 stage, meaning early segmentation occurs in a pre-cellular environment (Supplemental Fig S1). Frequency doubling of stripes of *Nv-slp* and *Nv-eve* RNA expression commences at the eve8 stage and proceeds in anterior to posterior progression, being finished in the 5th eve stripe by the wg7 stage (Rosenberg et al., 2014, Fig 1). The segment polarity genes *Nv-wg Nv-en*, and *Nv-prd* likewise express RNA in stripes that form in an anterior to posterior progression, and are only expressed in the ectoderm (as demarcated by *Nv-sim* expression), meaning that only the ectoderm autonomously segments in *Nasonia* (Fig 1). Together, these data support the notion that segmentation in the anterior proceeds in a progressive fashion, though the expression of *Nv-eve* and *Nv-runt* are exceptions to this.

**Figure 1:**
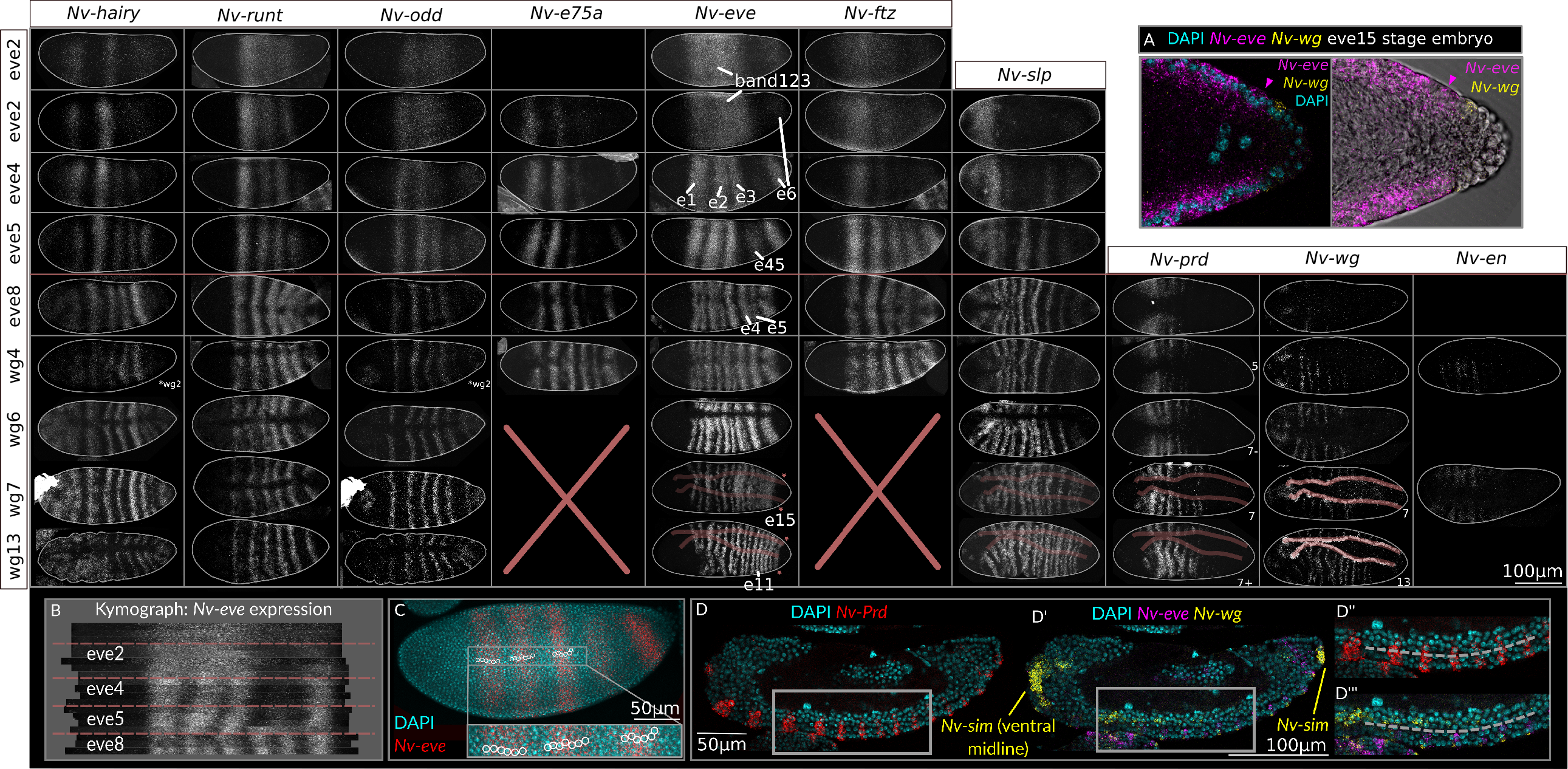
Pair rule gene expression during *Nasonia* development. All embryos are maximum intensity projections (half or full embryos), oriented anterior left and dorsal up. Embryos are laterally oriented unless indicated otherwise (D = dorsally oriented, V = ventrally oriented), and staged by the number of *Nv-eve* or *wg* stripes. Some embryos are re-used eg. *odd* and *hairy* expression comes from the same embryo. A: Magnified expression of the 15th, sequential *Nv-eve* stripe, note the single layer of cells around this stripe. B: kymograph of *Nv-eve* expression throughout segmentation. Each row represents expression of *Nv-eve* along the midline of the embryo, from the anterior to the posterior, and straightened in Image J. To account for variation in how the embryo midline is measured and embryo size, rows are aligned by the anterior of the first *eve* stripe and the posterior of the final *eve* stripe. Red dotted lines indicate the boundary of embryo stages. C: maximum intensity projection of an eve5 embryo. White lines surround cell nuclei to demonstrate that each *eve* stripe is six cells wide. D: *Nasonia* segment polarity genes are expressed in a one-cell repeat, as in *Drosophila* and other species. Images are single slices at the middle of the embryo. The boundary between the segmenting ectoderm and non-segmenting mesoderm is marked by a dashed line. Insets (D”, D”’) are taken from the region of the embryo marked by a thick white line.

The end of the embryo, 85-90% egg Length (EL), behaves differently to the rest of the embryo. *Nv-e75A* is expressed within this region until at least the wg4 stage (we did not image *Nv-e75A* stained embryos later than this), but other genes are not expressed in this stripe. The first segmental expression to be detected in this region of the embryo is that of *Nv-hairy* RNA, at the wg6 stage (Fig 1). Shortly after this, the 11th *Nv-eve*RNA stripe emerges from the anterior of the 6th *eve* RNA stripe (Rosenberg et al., 2014, Fig 1). Faint segmental expression of *Nv-slp* RNA is then observed, followed by splitting of *eve* RNA stripe six into three stripes of single segment periodicity, separated by a stripe of *Nv-slp* RNA expression (Fig S3). A sixth *Nv-odd* RNA stripe is detectable at this stage, but expression is delayed and very faint (Fig S3).

The posterior regions of *Nasonia* embryos segment sequentially. The first expression of pair rule gene RNA in this region is in a posterior cap of *Nv-odd* RNA, present from the eve4 stage. *Nv-hairy* RNA is expressed faintly and inconsistently the eve5 stage. Stripes of *Nv-ftz* and *Nv-runt* RNA appear posterior to the sixth eve stripe. Later, at the wg4 stage, the posterior *Nv-hairy* RNA expression becomes stronger and *Nv-odd* RNA is excluded from the *Nv-hairy* expression domain. After this, the fifteenth *eve* stripe appears in the posterior, appearing considerably thinner than the other stripes. This is the only sequentially appearing *Nv-eve* RNA stripe we observed, though Rosenberg et al., 2014 observe a sixteenth sequential stripe later. The sixteen stripes of the *Nasonia* are then established. This fifteenth stripe of *eve* RNA expression appears in the posterior of the *Nasonia* embryo before this region of the embryo has completed gastrulation (Fig 1A). This means that *Nasonia* is a long germband insect, patterning all segments before gastrulation (Davis & Patel, 2002).

We also observed that *Nasonia Nv-eve* RNA stripes were 5-7 nuclei wide, in contrast to *Drosophila* where pair rule RNA stripes are 3-4 nuclei wide (Schroeder et al., 2011). This doubling in size of the primary pair rule stripes does not correspond to an increase in the final segment polarity pattern: mature *Nv-wg* RNA stripes are one cell wide and separated by three nuclei (Fig 1B-D). How and why the pair rule pattern shrinks is unclear.

Though these distinct regions of the embryo differ in gene expression dynamics, there are some similarities between them. In all regions of the embryo, *Nv-hairy* is the first gene we investigated to be segmentally expressed, implying that *Nv-hairy* may play a key role in providing positional information to the *Nasonia* pair rule network. Secondly, *Nv-slp* segmental expression precedes *Nv-eve* stripe splitting.

### 2.2 *Nasonia* timer gene expression can produce progressive patterning

We wished to explain *Nasonia*’s progressive patterning, and the expanded initial pair rule pattern. We first stained for the proposed regulators of different phases of segmentation (*cad*, *D*, and *opa*) (Clark & Akam, 2016; Clark, 2017; Clark & Peel, 2018), and used these patterns as inputs to the model of pair rule patterning.

In the first five *eve* RNA stripes, the expression of *Nv-cad* and *Nv-opa* RNA complement each other: *Nv-cad* RNA retracts across the anterior-posterior axis, as *Nv-opa* RNA expands (Fig 2A-F). *Nv-cad* RNA retracts from the broad *eve* stripe (band123, 30-65%EL) region before segmental expression of *Nv-eve*, and continues to retract towards the posterior as *eve* stripe45 splits (Fig 2A,B, S5). *Nv-opa* RNA is first detected in the head and stripe 1 at the eve4 stage, and expands posteriorly at the eve5 stage-shortly before *Nv-eve* stripe splitting (Fig 2E). *Nv-D* is expressed with different dynamics. At the eve4 stage, *Nv-D* is expressed in a broad band, from the anterior of stripe 1 to the anterior of stripe 6. This expression persists until shortly before *eve* frequency doubling; at this stage *Nv-D* expression is lost in stripes 1-3 in a pair-rule-like pattern strongly resembling *Dm-D* (Nambu & Nambu, 1996; Russell et al., 1996). *Nv-D* expression is retained around stripe4/5 until these stripes start to undergo frequency doubling. Curiously, *Nv-cad* and *Nv-D* are never expressed within stripe 6 (which has unusual *eve* dynamics); this stripe progresses straight to *Nv-opa* expression (Fig 2E-G).

**Figure 2:**
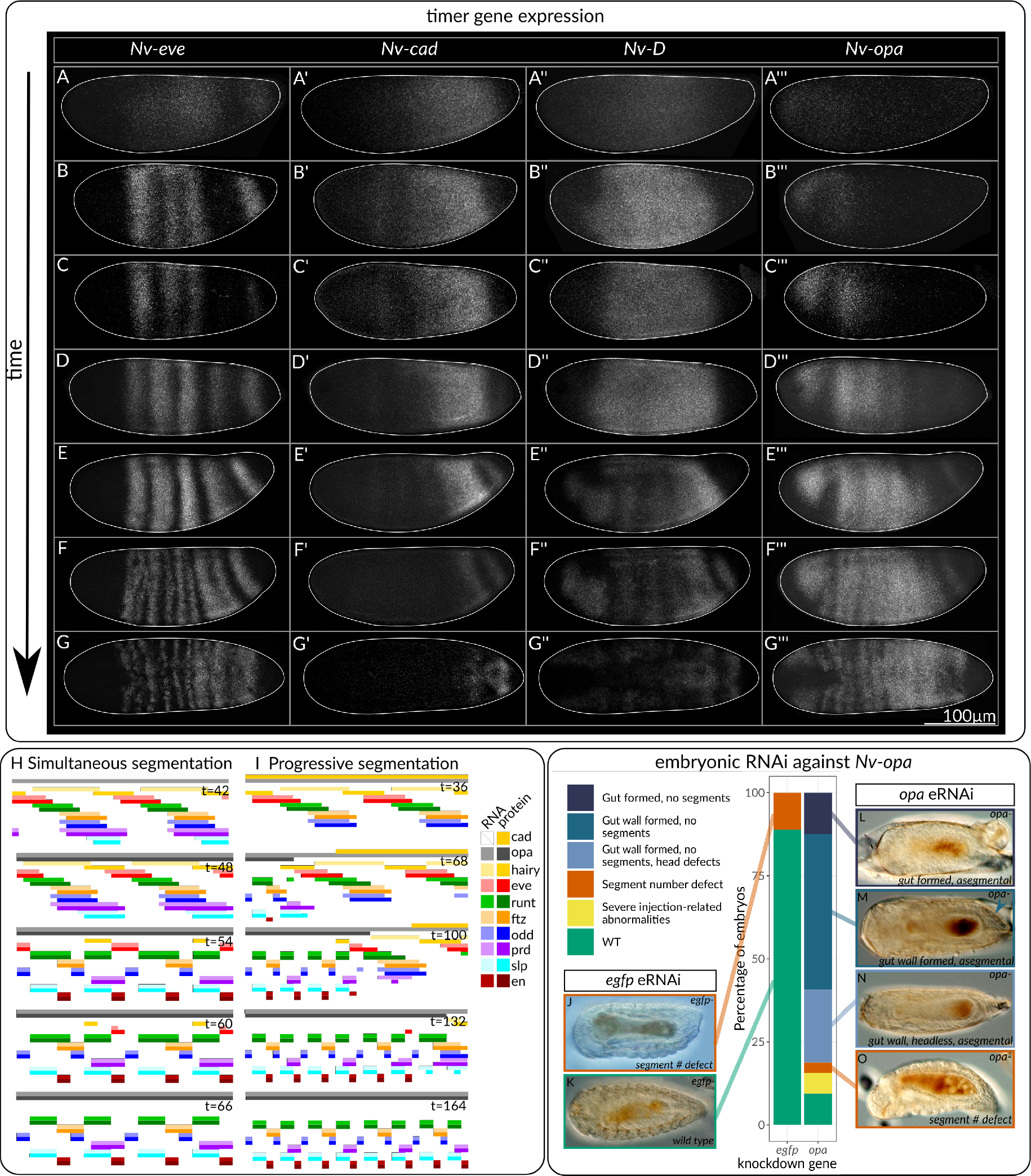
Timer gene expression in *Nasonia* recapitulates progressive patterning when modelled. A-G: Maximum intensity projections of embryos stained with the timer genes *Nv-cad, Nv-D, Nv-opa,* staged to *Nv-eve*. All embryos are laterally oriented, anterior left, except G which is ventrally oriented. H and I: simulation of broad pair rule stripes combined with simultaneous (H) and sequential/progressive (I) timer gene dynamics. J-O: *Nv-opa* is required for morphological segmentation. All embryos are oriented anterior left, ventral view. Arrow in M indicates the position of the gonads.

These *opa* and *cad* expression dynamics were able to recapitulate *Nasonia*-like progressive patterning, if simulated in a model containing *Nasonia*’s broad (6-cell wide) pair rule stripes (Fig 2H, I). In this model, stripes mature in an anterior to posterior progression, characteristic of progressive segmentation. The *cad* and *opa* dynamics are crucial to this progressive patterning: the same network simulated with broad pair rule stripes, but activated in a simultaneous manner results in an embryo that segments simultaneously, but has a final pattern doubled in size from the *Drosophila* model, ie a 16-cell repeat of gene expression (Fig 2H). The progressive model exhibits the *Drosophila* and *Nasonia*-like eight cell repeat. Note that the phasing of the late network in this simulation is *Drosophila*, not *Nasonia*, like.

To further investigate the role of the timer genes in *Nasonia* segmentation, we performed embryonic RNAi (eRNAi) against *Nv-opa*. We identified two phenotypes following *opa* eR-NAi: a total lack of segments within the embryo, and apparent defects in head formation. To distinguish between developmental arrest prior to morphological segmentation, and segmentation defects, we used DIC imaging to identify two morphological markers that appear after segmentation in wild type embryos: presence of the gut wall, and (where visible) gonads (Bull, 1982). Surviving embryos (8/67 *egfp*-, 32/64 *opa*-) were scored into six classes. Some embryos had an obvious gut, no gut wall, and no segments. These embryos were only present after injection with *opa* dsRNA, and could represent either developmental arrest at a stage prior to morphological segmentation, or a segmentation defect. Many embryos successfully formed the gut wall, which forms after morphological segmentation (Bull, 1982), but lacked any distinguishable morphological segments. Some of these embryos also lacked a head. A small number of *egfp* and *opa* injected embryos had defects in the number of segments. Control embryos with these phenotypes also exhibited cytoplasmic leakage, suggesting that this phenotype was caused by the injection procedure itself. Completely asegmental embryos occurred in 69% of surviving *opa-* embryos and never occurred in *egfp-* embryos. These data show that *Nv-opa* is required for morphological segmentation in *Nasonia*, supporting its proposed role as a key regulator of segmentation. The defects in head formation are consistent with the expression of *Nv-opa* in the head (Fig 2A-G), and its requirement for head formation in *Tribolium* (Clark & Peel, 2018).

We then attempted to understand how *Nasonia* undergo sequential segmentation / the appearance of stripe 15, again by staining for timer gene expression. Before stripe 15 appearance, at the eve9 stage, *Nv-cad* and *Nv-D* RNA are expressed in overlapping stripes behind the eve6 stripe, with *Nv-cad* most posterior and overlapping with *Nv-wg* RNA (Fig 3C). Later, at the eve9 stage, the fifteenth *eve* stripe appears within the *Nv-cad/Nv-D* domain, anterior to the posterior *Nv-wg* stripe (Fig 3C-F). At this stage, a posterior *Nv-opa* stripe is visible within the 6th *eve* stripe. This *Nv-opa* expression is stronger at the ventral ends of the embryo, and is expressed up to the anterior end of the fifteenth *eve* stripe (Fig 3D). At the eve15 stage, the *eve* stripe is still expressed within the *Nv-D* and *Nv-cad* domain, but *Nv-opa* has expanded posteriorly to be co-expressed with *eve* at the anterior of the *eve* stripe. Thus, *Nasonia* possess a similar spatial sequence of the timer genes as *Tribolium*: a spatial sequence of gene expression in the order *Nv-wg Nv-cad Nv-D Nv-opa*. This pattern was not simulated, because it is qualitatively similar to the sequential model of segmentation (Clark, 2017; Clark & Peel, 2018).

**Figure 3:**
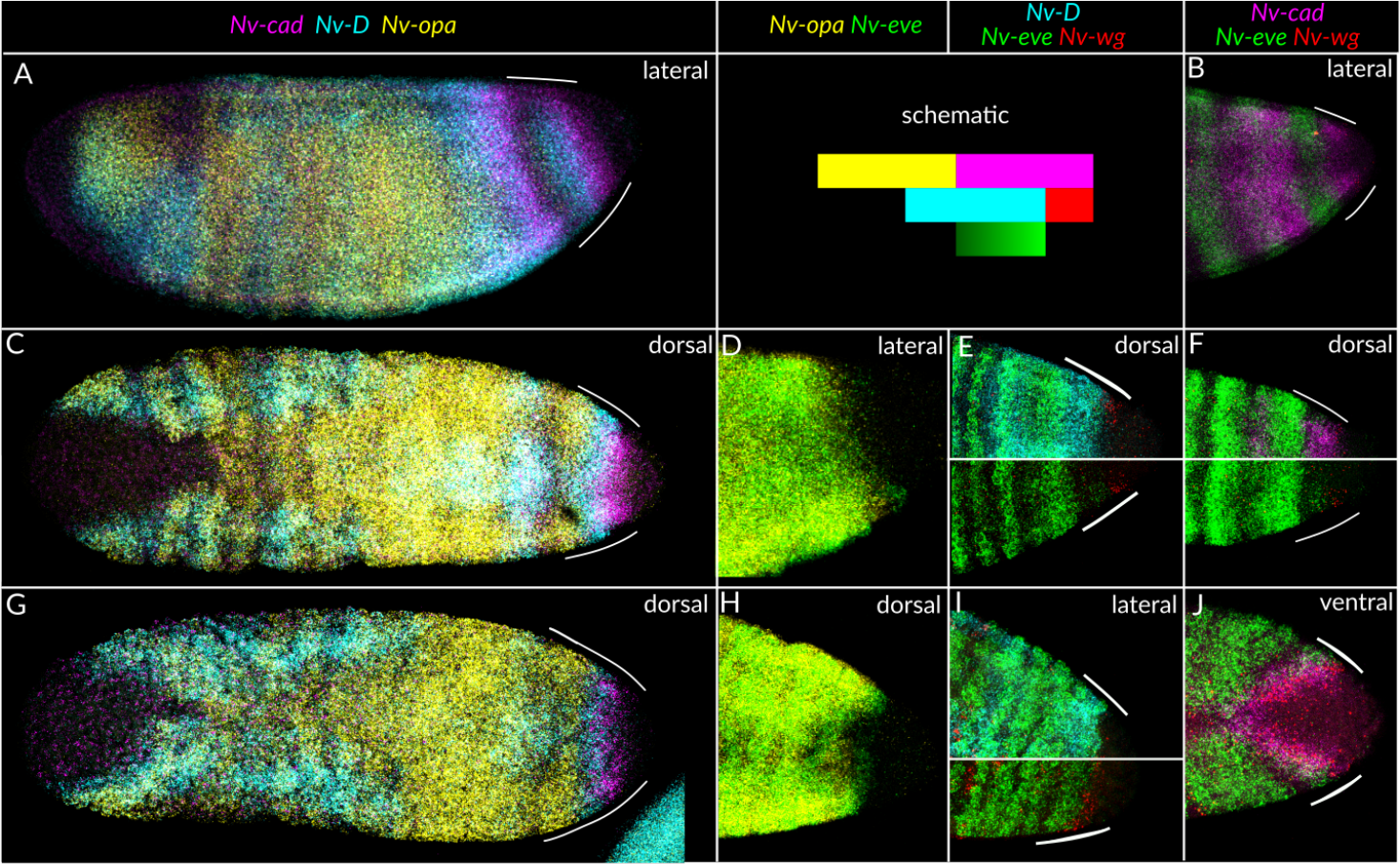
Timer gene expression in the posterior of the embryo is similar to other sequentially segmenting species. All embryos are partial or full maximum intensity projections, anterior left. Left panels (A, C, G): whole embryos. Right panels: cropped to just show the posterior.

With the exception of stripe 6, then, the *Drosophila* GRN combined with the *Nasonia cad* and *opa* expression patterns are able to recapitulate the *Nasonia* dynamics-both the existence of progressive segmentation in the anterior, and sequential segmentation in the posterior. However, this analysis relies on two critical assumptions. Firstly, it relies on there being two different pair rule GRNs in *Nasonia*, with qualitatively different behaviours-oscillatory vs non-oscillatory expression of genes. Secondly, it assumes that the pair rule networks of *Nasonia* and *Drosophila* are reasonably similar. In the following sections, we use our description of *Nasonia* segmentation to address both these points.

### 2.3 Organisation of *Nasonia* pair rule gene expression

Assessing whether two GRNs act in a given process is challenging. We observe, however, startling coordination between structural and behavioural changes in gene expression, which we interpret as meaning that two GRNs are acting in *Nasonia* expression. For instance, frequency doubling (of *Nv-slp* and *Nv-eve*) and expression of segment polarity genes begins at the eve8 stage, and occurs in an anterior to posterior progression within the embryo (Fig 1). Segmental expression of *Nv-prd* also begins at this stage, and again occurs in an anterior to posterior progression, with segmental expression of *Nv-prd* being detectable at the same time as *Nv-eve* and *Nv-slp* frequency doubling (Fig 1). There is also a dramatic shift in the relative expression of *Nv-slp* and *Nv-eve*: these genes go from being co-expressed to being strongly anti-correlated within the embryo, implying that the regulatory relationship between these genes has changed (Fig 5). These changes are tightly coordinated throughout the embryo, implying that they share a common cause. Expression of *Nv-opa* precedes these changes, implying that *Nv-opa* may cause these changes (Fig 2), an observation strengthened by the fact that *Nv-opa* is required for morphological segmentation. We also observe a change in dynamics at the eve8 stage. The second and third pair rule stripes shift forwards until the eve8 or eve9 stage, then stop (Fig 1B, Fig 4). This change in gene expression dynamics implies that the behaviour of the networks underpinning these gene expression patterns has changed. Together, these data imply that *Nasonia* possess two pair rule GRNs (or functional/pragmatic modules of a larger pair rule GRN (Verd et al., 2019)) with two distinct behaviours.

**Figure 4:**
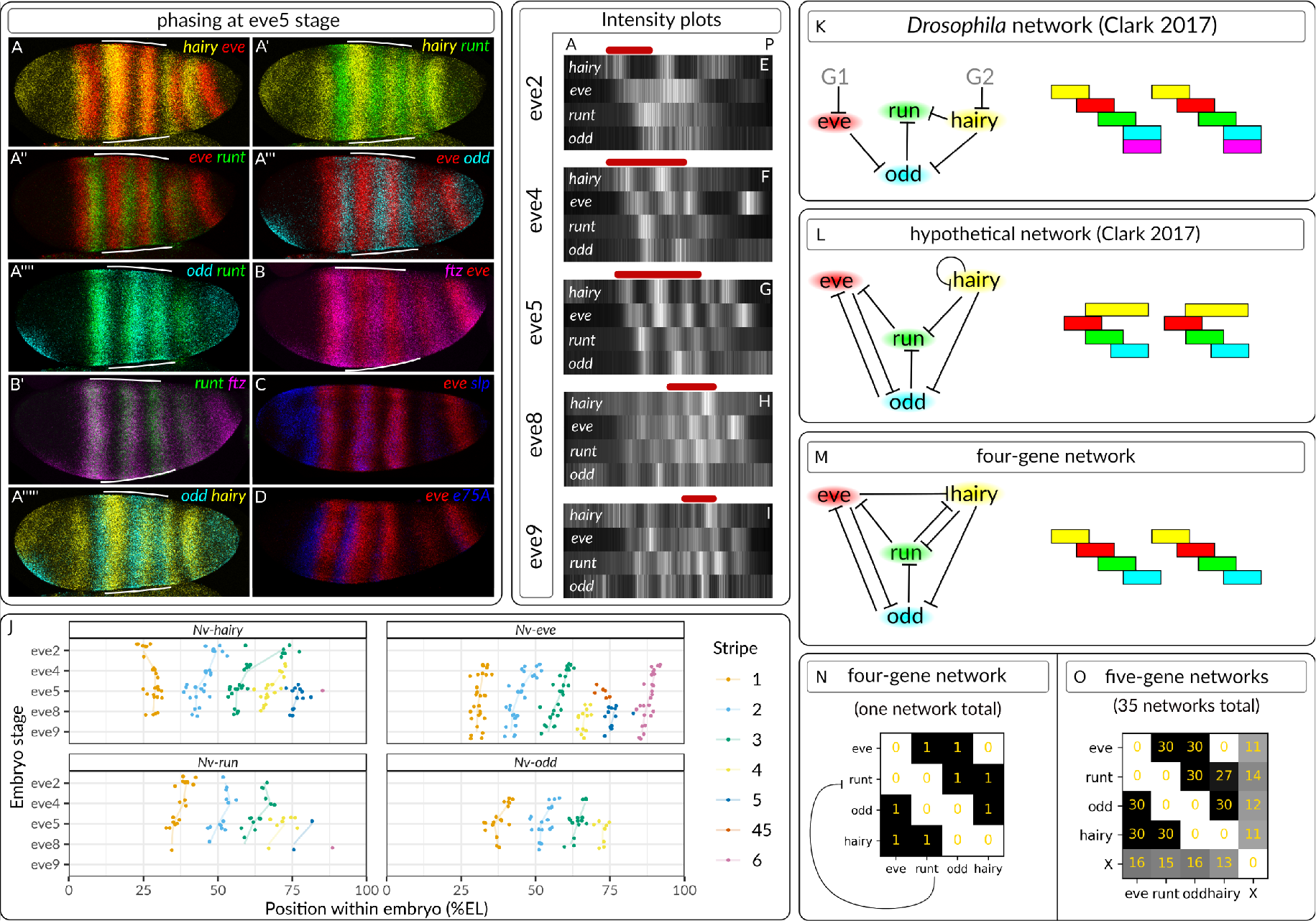
Phasing of the *Nasonia* early network closely resembles that of *Drosophila*. A-D: maximum intensity projections of embryos stained for pair rule genes. All embryos are laterally oriented, anterior left. White bars indicate the stripes which have the pair rule pattern established. E-I: Intensity plots describing gene expression at different embryo stages. Signal is averaged from 10-50*μ*m along a line following the curvature of the embryo, and are normalised 0 and 1. Background is defined as the signal intensity present in the head (where no genes except for *hairy* are expressed) and is set to zero. White bars indicate high intensity of gene expression. A: anterior. P: posterior. Red bars indicate the region of the embryo with the pair rule pattern established. Note that eve8 and eve9 embryos are beginning to undergo gastrulation so anterior stripes are more disordered. J: Forward movement of pair rule stripes within the embryo over time. Central positions of stripes were gauged by eye from straightened intensity profiles along the middle of the embryo. Black bars indicate standard error of the mean, coloured lines join the mean at different stages. K-M: network topology (left) and associated stable gene phasing (right). N: Topology of the four-gene network visualised as a matrix. Repressive interactions are read X to Y, that is, *run* represses *eve*. O: Five-gene networks possessing a given genetic interaction. Frequency of interaction is given in gold. Gene X: unconstrained fifth gene initialised with the same expression as *hairy*.

**Figure 5:**
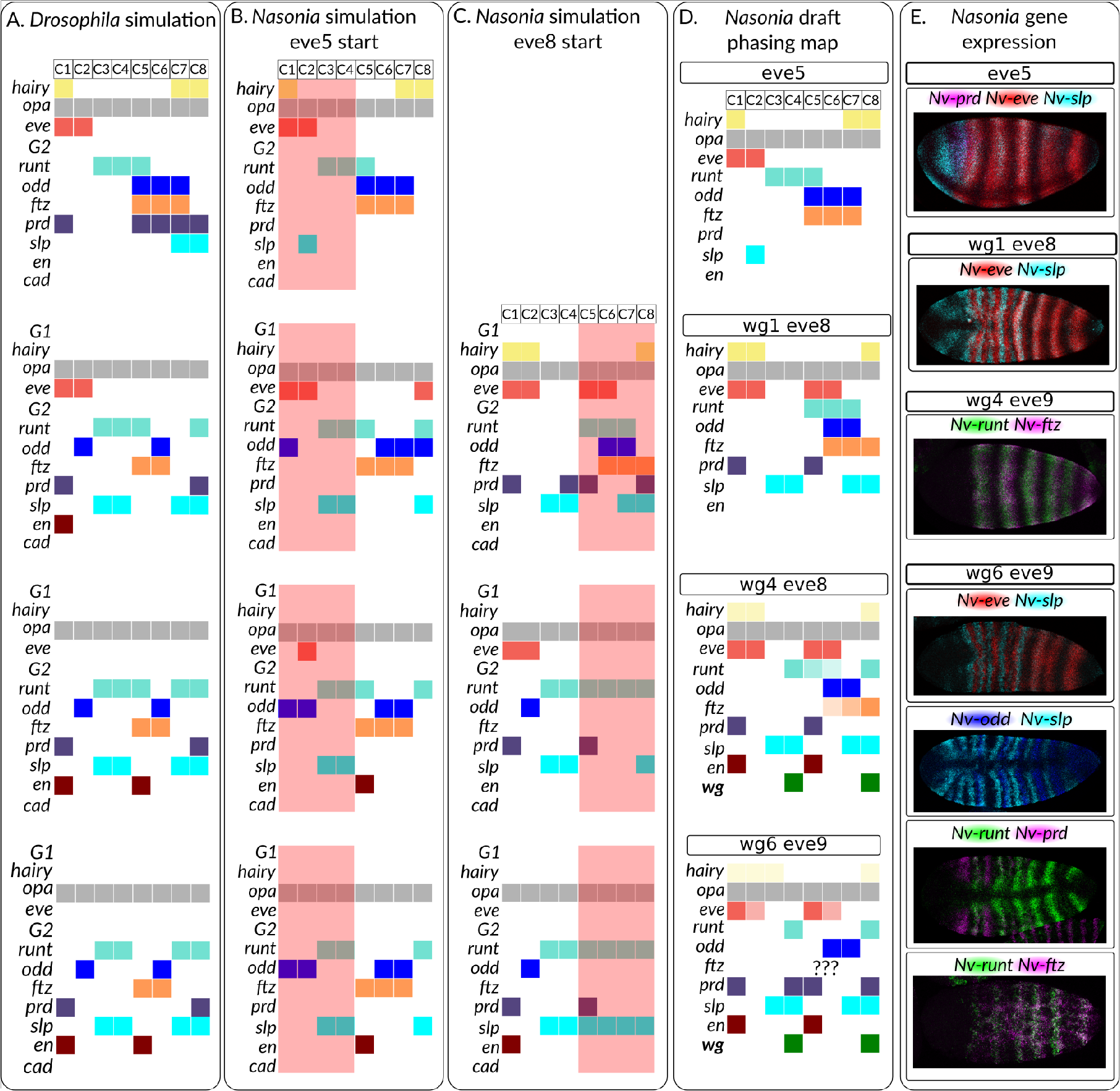
Altered inputs can recapitulate some aspects of late *Nasonia* segmentation. Note that each cell takes different numbers of time-steps to reach final output states so the temporal information depicted is not necessarily accurate. A: *Drosophila* simulations based on gene expression at the t=36 timepoint of simulations in Clark, 2017. Note that in cell 1 (C1), *slp* expression was omitted to ensure proper segmentation and because expression in this cell would decay without regulatory input. B: *Nasonia* simulations based on expression at the eve5 stage. Red bar indicates region where proper patterning is not produced, likely because of the lack of *prd* in the input. C: *Nasonia* simulations, starting with gene expression at the eve8 stage. Red bar indicates region where proper patterning is not produced. D: Draft map of gene expression in the *Nasonia* embryo. Map is based primarily of gene expression in the first four segments, which is representative of expression throughout the embryo. The map is produced based on comparisons to *Nv-eve* and *Nv-wg,* so other gene-gene comparisons may not be strictly accurate. E: selected HCRs showing relative expression of genes. All embryos are maximum intensity projections, anterior left. See Figs S6, S7, S8 for the rest of the dataset.

**Figure 6:**
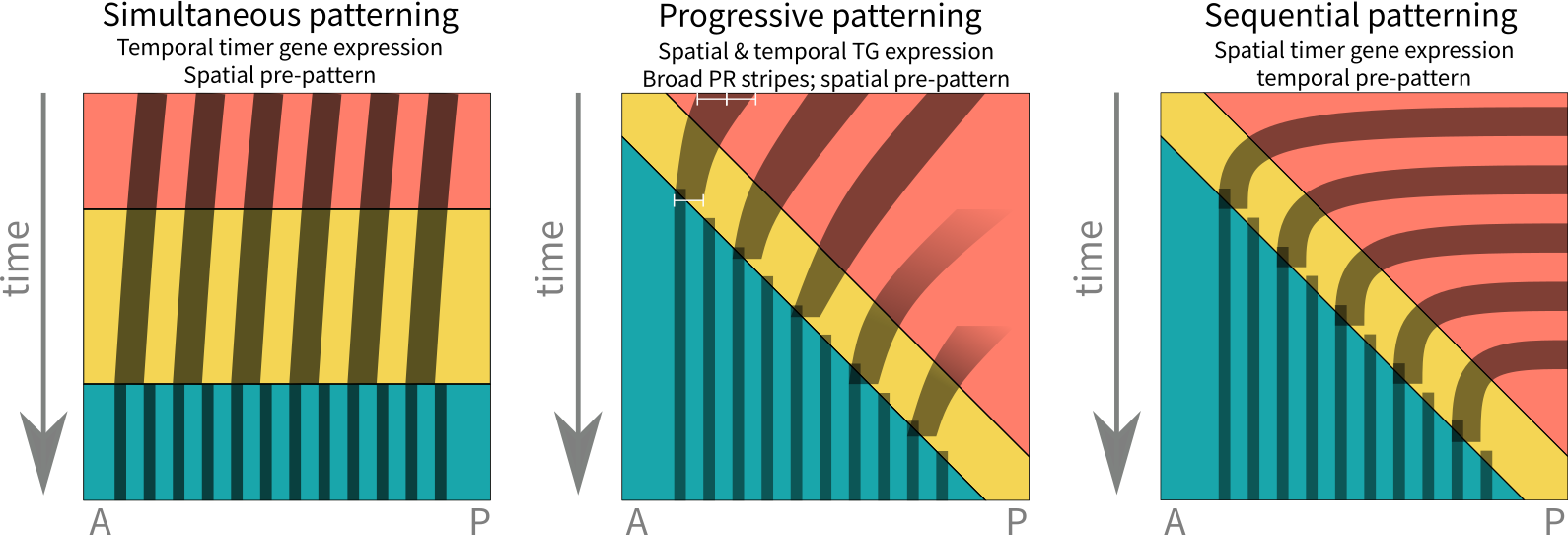
Schematic of simultaneous, sequential, and progressive segmentation. Adapted from Clark et al., 2019

### 2.4 Conservation of the early segmentation network

We then investigated the conservation of these two pair rule networks. The primary pair rule genes involved in the early network, *Nv-eve*, *Nv-hairy*, *Nv-odd*, *Nv-runt*, and *Nv-ftz*, are expressed in very similar patterns relative to each other as in *Drosophila* (Fig 4A-D, Rosenberg et al., 2014). *Nasonia* pair rule RNA stripes still overlap and are expressed in the order *Nv-hairy* → *Nv-eve* → *Nv-odd/Nv-ftz* → *Nv-runt* → *Nv-hairy* (see Fig 4A-D). To concisely display pattern maturation over time, we quantified gene expression along the midline of the embryo in ImageJ (averaging across 20-50*μ*m), and plotted these intensities as colour gradients. This analysis revealed that the *hairy-eve-runt-odd* pattern develops over time, from being present only in the very anterior at the eve2 stage, to being present in the fourth and fifth stripes at the eve9 stage (Fig 4E-I). The expression of *Nv-slp* and *Nv-prd* RNA, as mentioned above, has changed compared to *Drosophila*. *Nv-slp* RNA is expressed within, not between, *Nv-eve* RNA stripes (Fig 4C), and *Nv-Prd* RNA is not expressed until later in segmentation (Fig 1). In addition, we identified early segmental expression of *Nv-E75a* RNA anterior to *eve* RNA (Fig 4D), implying that this gene may also be involved in early segmentation.

These changes in *Nv-slp* and *Nv-prd* expression would not impact the behaviour of the early network, as these genes do not cross-regulate other pair rule genes (Clark, 2017). Expression of extra gene/s, eg *Nv-e75a*, is addressed below. Thus, the *Nasonia* expression of these genes is consistent with a topology very similar or identical to the *Drosophila* ones, and does not imply a change in behaviour of the *Nasonia* network.

Despite the above, we know that the *Nasonia* network is capable of different behaviour to that of *Drosophila*: sequential patterning. This would rely on oscillatory expression of the primary pair rule genes. We therefore looked for evidence that *Nasonia* have oscillatory pair rule gene expression, where oscillatory gene expression is expression of the full *hairy*-*eve-runt-odd* sequence within a single cell. We found no oscillatory expression in the anterior or posterior of *Nasonia*. In the posterior, sequential expression begins with a cap of *Nv-odd* expression (Rosenberg et al., 2014, Fig 1). At the eve8 stage, a posterior stripe of *Nv-runt* forms anterior to this *Nv-odd* cap. A stripe of *Nv-hairy* then subdivides the *Nv-odd* cap, and the fifteenth *eve* stripe is expressed. This produces a spatial pair rule gene sequence (anterior to posterior, *Nv-runt, Nv-odd, Nv-hairy, Nv-eve,* as well as a temporal *Nv-odd-Nv-hairy-Nv-eve* sequence in the region of the fifteenth eve stripe, again consistent with the sequence from *Drosophila*. However, no cell goes though the full gene sequence.

In the anterior, we were unable to determine gene expression in each cell over time, because of the shifting domains of gene expression and the lack of cell markers. We instead quantified the changes in stripe position over time, reasoning that changes in gene expression within a single cell would also be visible as changes in the position of stripes of pair rule gene expression within the embryo. Cells in the *Nasonia* anterior also do not go through a full cycle of gene expression, because though pair rule stripes shift forwards in the embryo, no pair rule genes move forwards by an entire segment. That is, no position within the embryo completes a full cycle of pair rule gene expression, starting and ending at the same gene. If this behaviour was due to stability, we would expect gene expression to arrest at one pair rule gene. It does not: all four pair rule genes where we had sufficient data to assess stripe shifts exhibit forward shifts in gene expression (Fig 4J). Thus, the *Nasonia* early network appears to be capable of oscillatory gene expression, though whether this is driven by the anterior movement of gap gene expression in *Nasonia*, or the intrinsic dynamics of the network, is unclear (Pultz et al., 2005; Brent et al., 2007; Clark, 2017; Ray et al., 2019). It is also worth pointing out that oscillations that do not go through a full cycle of gene expression (damped oscillations) can pattern segments in the gap gene system of *Drosophila* (Verd et al., 2018).

The early network topologies of Clark, 2017 are either incapable of sequential segmentation (the *Drosophila* network, Fig 4K), or inconsistent with the observed gene expression patterns in *Nasonia* (the hypothetical network, Fig 4L). To identify a four-gene topology capable of sequential segmentation and the *Nasonia* phasings, we performed a computational screen. Topologies were classed as potentially sequential if, after 100 time-points of the simulation (about 2.5 full oscillations), they went through the sequence *hairy → hairy/eve →eve → eve/odd → odd → odd/runt → runt → runt/hairy → hairy*. This screen identified one topology capable of producing sequential segmentation from normal inputs-the topology in Fig 4M. This shows that the gene phasing observed in *Nasonia* is consistent with at least one GRN topology capable of sequential segmentation. Unlike the hypothetical network, this potential network requires the positions of every pair rule gene to be provided to the simulation in order to produce normal patterning (Figure S4), meaning that simultaneous patterning using this topology requires an extensive (presumably gap gene-mediated) spatial pre-pattern. This topology also does not resolve the potential use of *Nv-e75a* and other novel pair rule genes, which could change the behaviour of the network. Accordingly, we screened for five-gene topologies capable of sequential segmentation, with the fifth gene (gene X) initially co-expressed with *hairy* but with no final restrictions on its expression. After filtering out networks with topologies identical to the four-gene network, this analysis identified 35 potential networks, for which the frequency of genetic interactions are presented in Fig 4O. Although some genetic interactions remained impossible for this model formulation, the addition of the extra gene provided flexibility to the network, ensuring that no genetic interaction was present in every predicted network. Though this analysis is unlikely to have identified every possible five-gene network capable of sequential segmentation, as the initial conditions of gene X could be very different to our assumptions, it does demonstrate that there are various possible networks incorporating different genes (eg *Nv-e75A*). We draw two conclusions from the above analysis. Firstly, we identified one potential network capable of producing both the *Nasonia* gene expression patterns and sequential patterning. Secondly, addition of one extra gene (plausibly *Nv-e75a*) provides more flexibility to genetic interactions in the early network but does not have to change the behaviour of the network.

Taken together, this analysis shows that the *Nasonia* network could be capable of oscillatory gene expression, and sequential segmentation.

### 2.5 Topological changes required to explain some aspects of *Nasonia* late network

Unlike the early network, there are changes to gene expression in the *Nasonia* late network. Different genes undergo frequency doubling in *Nasonia* compared to *Drosophila*. In *Nasonia*, only *Nv-eve*, *Nv-slp*, and *Nv-runt* undergo frequency doubling, whereas in *Drosophila*, *Dm-odd* also doubles (Schroeder et al., 2011; Rosenberg et al., 2014). *Nv-odd* is expressed as a two-cell wide stripe, not one-cell wide, while secondary *Nv-runt* stripes are expressed very briefly and are only one cell wide (Fig 1, Rosenberg et al., 2014). In addition, *Nv-slp* and *Nv-odd* overlap (Fig 5E). We wished to know whether the observed changes in expression in the late network could be explained by changes in input to the late network-ie the altered positioning of *Nv-slp*, *Nv-prd*, and *Nv-E75A* or whether they require changes in gene regulation.

We approached this problem via simulation. We used an R package, BoolNet (Müssel et al., 2010), to model the output of the *Drosophila* network under various input conditions. This package ignores the gap between RNA transcription and protein translation, unlike Clark, 2017’s model. We used this abstraction to avoid arbitrarily assigning protein and RNA ages when initialising models. Modelling *Drosophila* gene expression at the t=36 timepoint in this way produces gene expression patterns identical to the existing model if *slp* expression in cell 1 is ignored (this expression would decay naturally in the next few timepoints) (Fig 5A). Modelling the *Nasonia* eve5 gene expression was able to produce some aspects of *Nasonia-* like patterning of the even-numbered segment. In this segment, the main difference between *Drosophila* and *Nasonia* is that *odd* stripes are two cells wide, and that these cells are co-expressed with *slp* (Fig 5D). Additionally, towards the end of segmentation, *Nv-runt* is expressed in one-cell wide stripes, co-expressed with *Nv-wg*. The altered inputs to the even-numbered parasegments are able to recapitulate these patterns: the lack of *slp* expression posterior to *odd* means that the *odd* and *ftz* domains are not repressed posteriorly, and these genes remain as two/three cell stripes. In these simulated cells, *slp* and *prd* expression do not resolve properly-*slp* because, in *Nasonia*, *slp* and *odd* are co-expressed, a pattern inconsistent with the mutual repression of these genes in *Drosophila* and the *Drosophila* model used. *Prd* never becomes expressed because *prd* expression is only possible in cells already expressing *prd* and we supply no *Nv-prd* expression to the model. As in the *Drosophila* model, secondary *eve* stripes do not appear. In summary, this analysis shows that the *Nasonia* inputs to the *Drosophila* network are able to produce some of the altered *Nasonia* gene expression.

We were unable to produce proper patterning in odd-numbered segments. Second *runt* and *odd* stripes form, which are not present in *Nasonia* embryos, and *odd* is mis-positioned.

To check that this was not due to the lack of *Prd*, we initialized the model with *Nasonia* inputs from the eve8 stage. This produced *Drosophila*, not *Nasonia*, gene expression patterns, and incorrect phasings in C4-C8 (compare Fig 5A-C). This means that the *Nasonia* inputs, at both the eve5 and eve8 stage, cannot produce the *Nasonia* outputs, and therefore that the *Drosophila* model cannot produce the *Nasonia* odd-numbered stripe gene expression. Involvement of an additional gene is required to produce these changes. In the *Nasonia* cell expressing *odd*, *eve* is the only gene to be expressed, both in simulations and the embryo (Fig 5B-D). Eve is thus the only modelled pair rule gene that could turn *odd* off in this cell. However, we know that a repressive relationship between *odd* and *eve* cannot exist in *Nasonia,* because these genes are stably co-expressed (Fig 5D). Thus, additional genes must be involved to repress *odd* in this cell. A similar argument holds for *runt*: in cells 3-4, only *slp* is co-expressed with *runt* so only *slp* could repress *runt*. Again, there cannot be a repressive interaction between these genes, because *slp* is co-expressed with *runt* in cell 8.

This means that the *Drosophila* network can produce *Nasonia*’s expanded *odd* and *ftz* expression, as well as the smaller *runt* domains. However it cannot recapitulate the lack of *odd* doubling, and general patterning in cells 1-4. Changes in gene regulation possibly related to e75a and/or other genes are required to explain these features.

## 3 Discussion

Here, we explore the conservation of the pair rule GRN in *Nasonia* and *Drosophila*. Despite morphological and gene expression differences between *Nasonia* and *Drosophila* segmentation, we found that altered inputs to a largely conserved *Drosophila* network can explain these differences. The general organization of the pair rule genes into two networks is likely conserved in *Nasonia*, providing the first evidence of such organization outside *Drosophila*. In addition, though there are changes to the genes expressed in the early network, the early network still appears to behave in a similar way to *Drosophila*. We were able to recapitulate the *Nasonia* segmentation dynamics-progressive patterning-using the *Nasonia* timer gene expression patterns. Finally, changes in the input to the late network are able to recapitulate some, but not all, changes in *Nasonia* segmentation; there must be some topological changes to the late network.

### 3.1 Timer gene hypothesis explains the dynamics of *Nasonia* segmentation

Clark, 2017 had previously shown that a slightly modified *Drosophila* pair rule GRN can perform both simultaneous and sequential patterning. That is, either spatial or temporal information can be used to produce patterning. In simultaneous patterning, the information is spatial, in the form of gap gene mediated patterning of pair rule genes. In sequentially segmenting insects, a temporal pattern is used (Clark, 2017; Clark & Peel, 2018; Clark et al., 2019). Our work reinforces and extends both these points. Firstly, we analyse an insect which combines multiple modes of segmentation (Rosenberg et al., 2014). In the anterior, *Nasonia* undergo progressive patterning, while in the posterior, they undergo sequential segmentation. Anterior expression of timer genes was used to model progressive patterning, while in the posterior, the timer genes are expressed in a similar spatial sequence to the other sequentially segmenting insect studied, *Tribolium* (Clark & Peel, 2018). Presumably, the GRN used does not change between the anterior and posterior of *Nasonia*. Together, these findings mean that the expression of timer genes can explain the unique segmentation dynamics of *Nasonia*.

Secondly, inspired by *Nasonia* patterning, we provide a way to combine a spatial and temporal pre-pattern, producing what we have called *progressive patterning*. Broad pair rule stripes (an expanded spatial pattern) correct for the temporal pattern, an expansion/retraction of *cad* and *opa* expression, ultimately leading to progressive patterning.

This finding has important implications for the evolution of segmentation. It means that full simultaneous and sequential segmentation are likely two ends of a spectrum, rather than distinct types of segmentation. Such a spectrum is implied in the findings of Clark, 2017; Clark and Peel, 2018: if simultaneous and sequential segmentation share a mechanistic basis or GRN, then intermediates could exist. Small changes to the expression of the timer genes, and the width of pair rule stripes, can have dramatic impacts on how the pair rule GRN behaves, and so to the dynamics of segmentation, providing a simple and elegant method of evolving phenotypic variation.

### 3.2 Topological changes to network

Unsurprisingly, modelling the dynamics of *Nasonia* segmentation was unable to recapitulate differences in gene phasings between *Nasonia* and *Drosophila*. Our data, combined with modelling, provide insight into potential regulatory changes achieving these altered expression patterns in *Nasonia*. We found that the general organisation of the *Drosophila* pair rule network was conserved in *Nasonia*, as apparent changes to gene regulation are coordinated within the embryo, and correlate with *Nv-opa* expression. This conservation of GRN structure implies that such a structure is evolutionarily important (Clark & Akam, 2016; Clark & Peel, 2018). It also provides support for the timer gene idea, which relies on two GRNs with different behaviours being activated in different ways, to achieve patterning (Clark, 2017).

We found no evidence for topological changes to the early network, aside from the changes in *Nv-slp* and *Nv-prd* expression. The primary pair rule genes are expressed in the same order as *Drosophila*, and like *Drosophila*, shift anteriorly within the embryo over time (Fig 4A-J, (Clark, 2017; Lim et al., 2018)), though whether this is driven by gap genes is unclear (Pultz et al., 2005; Brent et al., 2007; Ray et al., 2019). This means that the *Nasonia* early network can be modelled by the *Drosophila* GRN. We identified a four-gene network topology capable of sequential segmentation, meaning that the expression patterns observed are consistent with sequential segmentation. Finally, we found that addition of extra genes to this network does not change the behaviour of the network, meaning the possible addition of extra genes, eg. *Nv-e75a,* to the early network does not imply a change in network behaviour. This shows that the *Nasonia* early network is similar in behaviour to the *Drosophila* one, and that such behaviour is consistent with a network capable of sequential segmentation.

We also found that the *Drosophila* late network can explain some, but not all, aspects of *Nasonia* segmentation. The expanded *odd* and *ftz* domains in *Nasonia* can be explained by the *Drosophila* network and *Nasonia*’s altered inputs to it, while the lack of *odd* frequency doubling requires topological changes to the network. We are able to predict some of these changes. Firstly, *Nasonia* must lack strong mutual repression between *slp* and *odd* to maintain stable co-expression of these genes. Secondly, there must be another gene or genes patterning the anterior parasegment/cells 1-4, to prevent frequency doubling of *Nv-runt* and *Nv-odd*. Thirdly, the regulation of *Nv-eve* must change to allow *eve* frequency doubling. Crucially, there is no evidence that such changes would change the behaviour of the network-the *Nasonia* late network still has a similar end point: stable expression of segment polarity genes (Figs 1, S2). This is consistent with previous work, as empirical and modelling work imply that the segment polarity network is stable and well conserved (Von Dassow & Meir, 2004; Choe & Brown, 2009; Janssen & Budd, 2013; Green & Akam, 2013; Vellutini & Hejnol, 2016; Auman & Chipman, 2018; Clark et al., 2019).

### 3.3 Studying GRNs

These findings reinforce a number of important points. These behaviours of the *Drosophila* and *Nasonia* pair rule networks can only be understood using dynamical modelling ((Clark, 2017), Figs 4, 2), reinforcing the time-dependent nature of the genetic regulation of development, and the necessity of using modelling as a tool to understand GRNs (DiFrisco & Jaeger, 2019; Briscoe, 2019). Secondly, the late GRN is input-dependent: changes to the inputs of this GRN are able to explain some (but not all) changes in the *Nasonia* gene phasings of the late network. Again, this reinforces the need to use modelling alongside empirical research, and in particular the power of using modelling to understand developmental processes in less-studied species such as *Nasonia*.

We also found that though the dynamics of *Nasonia* patterning can be recapitulated using the *Drosophila* network, the relative patterns of gene expression cannot be fully accounted for by such an analysis.

Are the *Nasonia* and *Drosophila* GRNs homologous? They share common descent: the involvement of pair rule genes in arthropod segmentation is extensively documented, and is at least as old as holometabolous insects (Dearden et al., 2006). It is unclear whether these GRNs are structurally homologous, as we collected no data on the interactions between *Nasonia* pair rule genes. Moreover, the structure of a GRN provides little information as to the behaviour of such a GRN within a cell, and what traits it can allow in an organism (DiFrisco & Jaeger, 2019). However, we do provide evidence that the two GRNs are functionally homologous, that is, they behave the same way within the organism. The early network, as argued above, looks very similar to *Drosophila*. The late network is more difficult: there appear to be some changes to gene regulation. However, the output of segmentation is the same (expression of the segment polarity genes), and there is no evidence for a change in the dynamics of stripe expression (eg anterior movement of stripes). We think that this conservation of output and behaviour means that the function of the late network is unchanged. Thus, the *Drosophila* and *Nasonia* GRNs may be thought of as functionally homologous (DiFrisco & Jaeger, 2021).

### 3.4 Conclusion

Overall, we find that remarkably few changes to the *Drosophila* pair rule GRN are required to simulate *Nasonia* patterning. *Nasonia*’s progressive patterning can be recapitulated by changing the *Nv-cad* and *Nv-opa* dynamics, while some changes to the late network can be simulated using only the changed *slp* and *prd* expression at the eve5 stage. Our method can provide no direct evidence that specific interactions in the *Drosophila* GRN are conserved in *Nasonia*, but this analysis does imply that if there are substantial topological changes to the *Nasonia* network, these do not result in changes to the patterning process.

Finally, the similarities between *Nasonia* and *Drosophila* segmentation, at the level of the GRN involved, implies that these derive from a common ancestor GRN that likely evolved deep in the insect lineage. This GRN has proven developmentally flexible over evolutionary time, allowing different forms of morphological segmentation to be built on an overall conservative network. The changes we have detected in the GRN that underlie *Nasonia* segmentation are limited implying that only minor modifications of an ancestral but flexible GRN may be enough to generate endless variety in morphological segmentation.

## 4 Methods

### 4.1 Hybridization chain reaction

*Nasonia* were raised on commercially sourced *Sarcophaga bullata* pupae (from Mantis Mayhem, https://mantismayhem.co.uk/shop/ols/products/green-blue-bottle-flies-pupae), at 25°C. *Nasonia* cultures were a kind gift from Dr David Shuker and Dr Nicola Cook (St Andrews University).

Embryo collection was carried out after Werren and Loehlin, 2009. Flugs were modified such that *S. bullata* pupae could be placed, head-exposed, inside the plug. Embryos were collected from pupae provided to *Nasonia* for 12 hours, to provide the relevant stages of segmentation. Some embryos were stored in the fridge overnight for convenience. Hosts were cracked open under a dissecting microscope and dipped into 5mL of heptane, 4.5mL of PBS (phosphate buffered saline)) and 0.5mL 37% formaldehyde in a 15mL falcon, and fixed for 8-18 hours. After fixing, embryos collected at the bottom of the falcon. To dechorionate the embryos, the bottom formaldehyde layer was replaced with 100% ice cold methanol, and shaken vigorously for 1-3 minutes. Dechorionated and devitellenized embryos settled at the bottom of the falcon and were transferred to an eppendorf, washed 3x in methanol, and stored at −20°C.

Hybridization chain reaction was performed per Choi et al., 2016. Proteinase K digestion was not required for efficient probe penetration, so embryos were dehydrated in a methanol series before incubation in 100 *μ*L hybridization buffer for 30 minutes at 37°C was added. 1 pmol probe (ie 1 *μ*L of 1*μ*M probe) was added to 50 *μ*L hybridization buffer, and embryos were incubated with probe overnight at 37°C. Embryos were then washed four times with probe wash buffer for 15 minutes each at 37°C, before being washed three times for five minutes each in 5 x SSCT (5 x sodium citrate-Tween buffer). Embryos were pre-amplified in amplification buffer for 30 minutes. Hairpins were prepared by heating each hairpin individually to 95°C for 90 seconds, and leaving in a drawer at room temperature for 30 minutes. 3 pmol hairpin was added to 50 *μ*L of amplification buffer and added to the embryos. Embryos were amplified overnight in the dark at room temperature. Embryos were then washed 2x 5 minutes, 2x 30 minutes, and 1x 5 minutes in SSCT. DAPI (Invitrogen) was added to the first 30 minute wash. Embryos were mounted in ProLong Glass (Invitrogen), left to cure at room temperature overnight, and stored in the fridge before imaging. Imaging was performed using a Olympus FV3000 confocal microscope at the Department of Zoology, Cambridge University, using the UPLSAPO30X 30X silicon oil lens.

### 4.2 Modelling

All scripts are available at https://github.com/Shannon-E-Taylor/masters.

Simulations of the pair rule system were performed using the modelling framework described in Clark, 2017, and the code from the supplemental information of that paper. The model consists of a one-dimensional row of cells. Boolean network analysis was performed using the BoolNet R (R version 3.4.4) package (R Core Team, 2018; Mussel et al., 2010). The full GRN from the supplemental information of Clark, 2017 was used for modelling. To generate a state graph, the `plotStateGraph` function was used. To identify attractors, the `getAttractors` function was used; the default version of this code identifies all attractors for a synchronous network using the `exhaustive` method, which identifies trajectories for every possible initial condition.

### 4.3 Image analysis

Image analysis was carried out using Fiji (Abràmoff et al., 2004; Schindelin et al., 2012). Analysis of blastoderm-stage embryos was done on partial or full maximum intensity projections; where embryos had more than one cell layer, relationships between genes were confirmed using the original 3D images. Background fluorescence was defined as the fluorescence visible in regions of the embryo clearly not expressing the gene of interest (for most embryos, the head) and was removed using the brightness-contrast tool in Fiji.

### 4.4 Embryonic RNA interference

Embryonic RNAi (eRNAi) was performed in Aotearoa New Zealand using a different strain of wasps to those in Cambridge. These *Nasonia* were reared on *Lucilia sericata* blowflies (www.biosuppliers.com), using similar methods as Werren and Loehlin, 2009 (*S. bullata* are not commercially available in Aotearoa).

dsRNA against *egfp* and *Nv-opa* was ordered from Genolution (http://genolution.co.kr/agrorna/service-overview/).

To prepare embryos for microinjection, adult wasps were exposed to hosts for 2-5 hours. This long exposure time led to embryos of very similar stages, because the *Nasonia* took several hours to prepare to lay. Embryos were gently collected using fine forceps and aligned on a 1% agarose/PBS plate. Embryos were affixed to coverslips using heptane glue and covered with a small drop of *Drosophila* microinjection oil (1.75mL Halocarbon oil 700 + 0.25mL Halocarbon oil 27, Sigma). For the experiment reported here, embryos were then left in the fridge for four hours until it was time to inject them, these embryos were blastoderm stage embryos prior to cellularization. Other experiments, injecting embryos at the time of pole cell formation, resulted in similar phenotypes.

Embryos were injected using the PLI-100 (Harvard Apparatus) injection apparatus, and borosilicate needles. The coverslip was then transferred to the agarose plate to maintain humidity, and incubated at 28°C until all embryos had fully developed, 24-36 hours. Imaging was performed using DIC optics and an Olympus BX61 compound microscope.

## Acknowledgements

We are very grateful to Prof. Michael Akam, Dr. Matthew Benton, Dr. Eric Clark, Prof. James Maclaurin, Dr. Olivia Tidswell and Dr. Berta Verd for very helpful advice and discussion. We would like to thank Petra Dearden for proof-reading this manuscript. We are also grateful to the Cambridge Advanced Imaging Centre for assistance with imaging. This work was part funded by Genomics Aotearoa.

