## Supplemental Figures for "*Nasonia* segmentation is regulated by an ancestral insect segmentation regulatory network also present in flies"

### 6 Supplementary Material

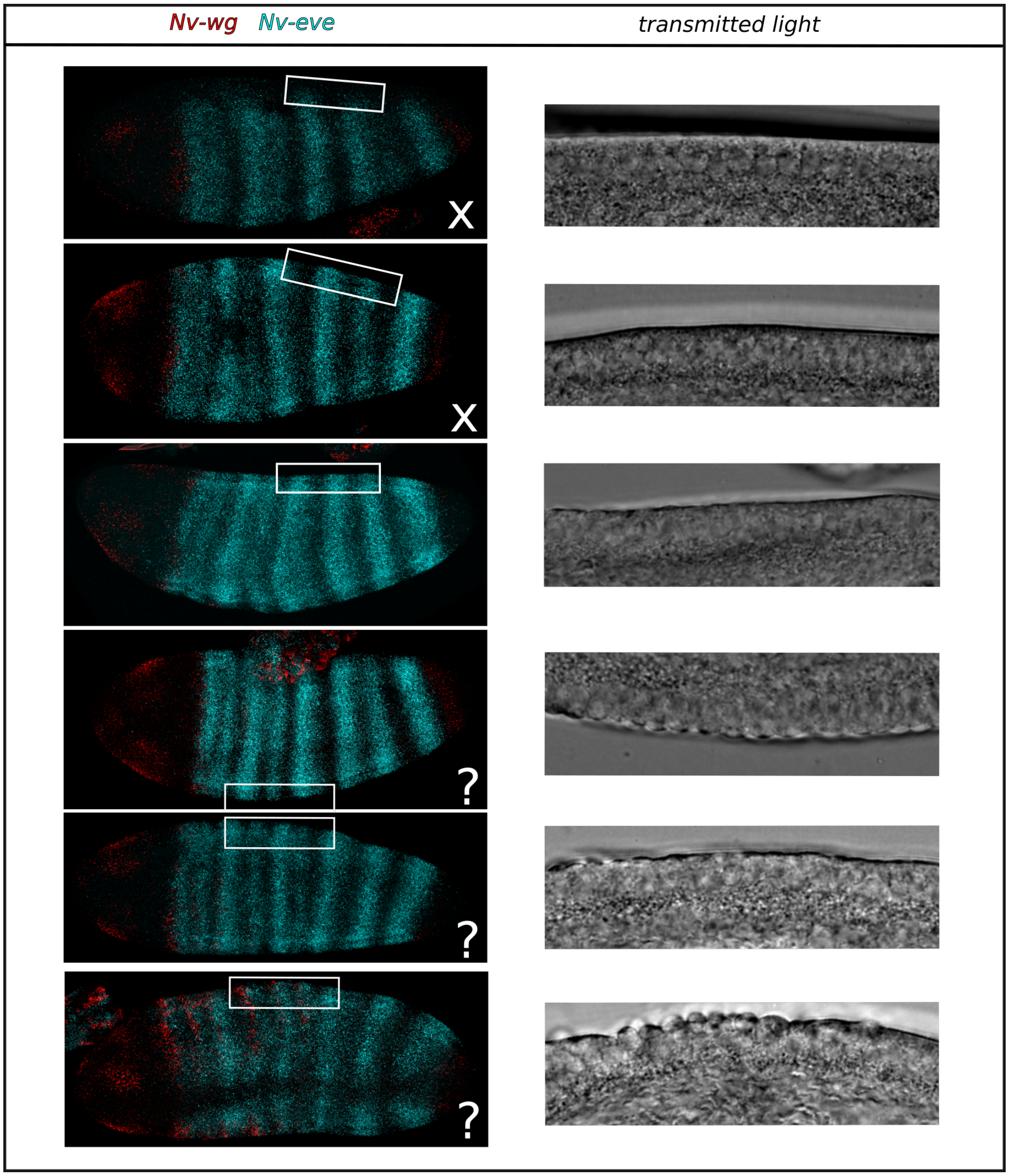


Figure S1: Timing of cellularization relative to segmentation in *Nasonia*. Left panel: maximum intensity projections of embryos stained for *Nv-eve* and *Nv-wg*. Anterior left. Right panel: enlargement of transmitted light of one optical section towards the middle of the embryo. Embryos are arranged in a temporal progression, younger embryos at the top of the page. X's indicate that an embryo is definitely not cellular; ?'s indicate embryos where cellularization has begun.

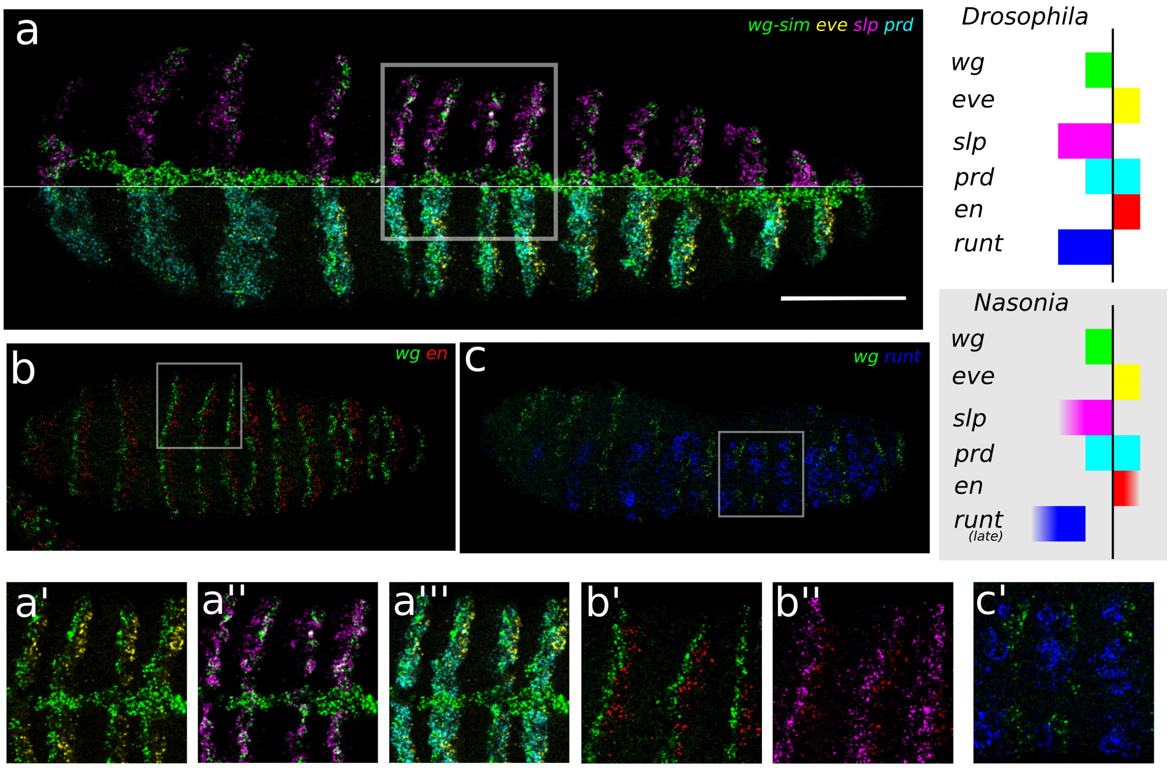


Figure S2: Gene expression across the segment polarity boundary of *Nasonia*. *Drosophila* schematic given is adapted from Green and Akam 2013; *Nasonia* schematic is inferred from the data. Gradients represents uncertain boundaries where genes abutting that boundary were not present in the sample stained. Vertical black line depicts the presumptive parasegment boundary. Scale bar: 50um. Boxes outline regions of the embryo enlarged in the bottom panel. All images are single confocal slices of representative germband extended embryos, ventral view. Horizontal white line in **a** marks where the genes depicted change in this embryo. **a'**-**a"'**, **b'**-**b"**, **c'**: enlargements of indicated regions of the embryos. Note that the *Nv-runt* expression has faded then re-emerged.

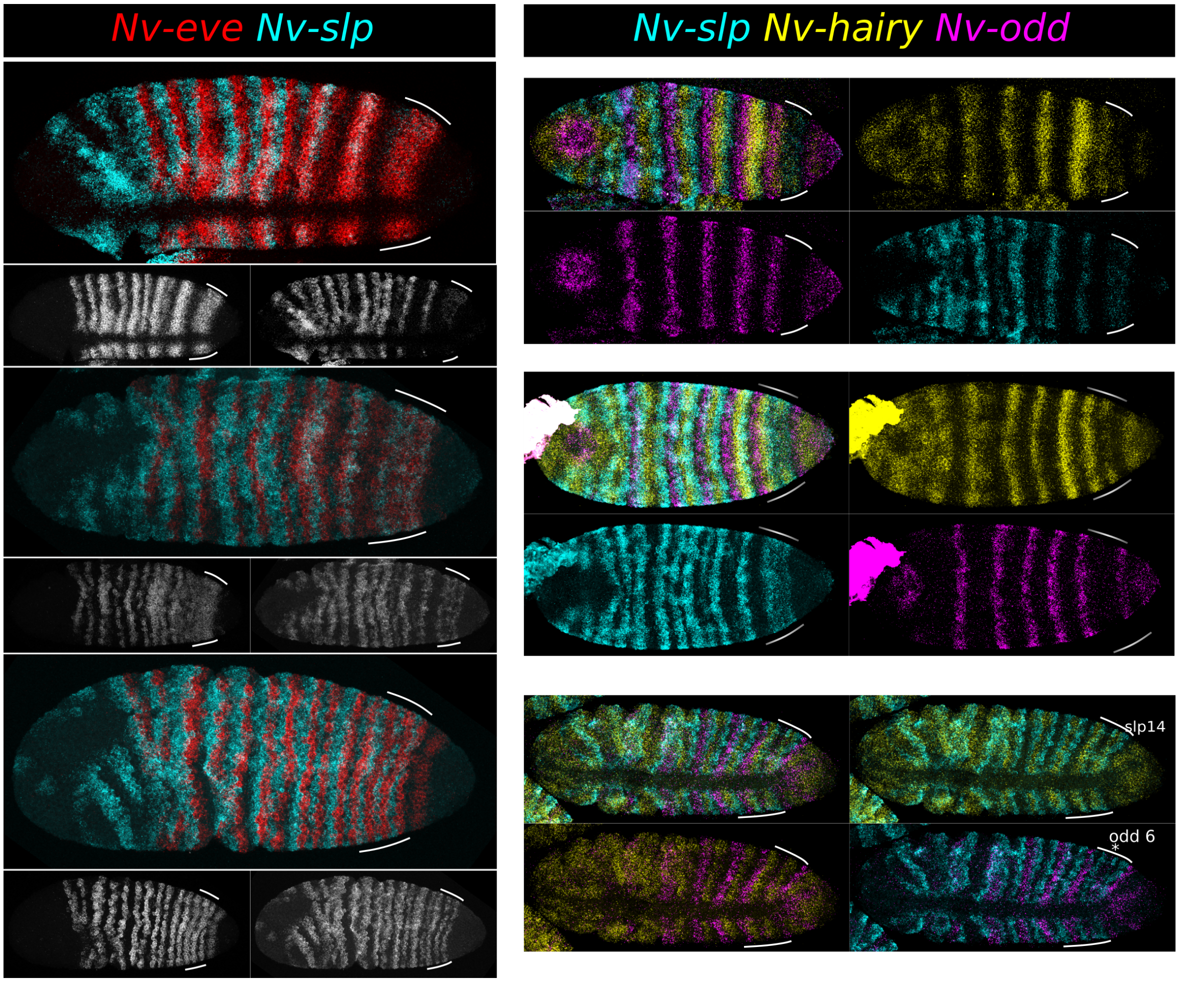


Figure S3: Gene expression within the e6 stripe. All embryos are maximum intensity projections, laterally or dorsally oriented.

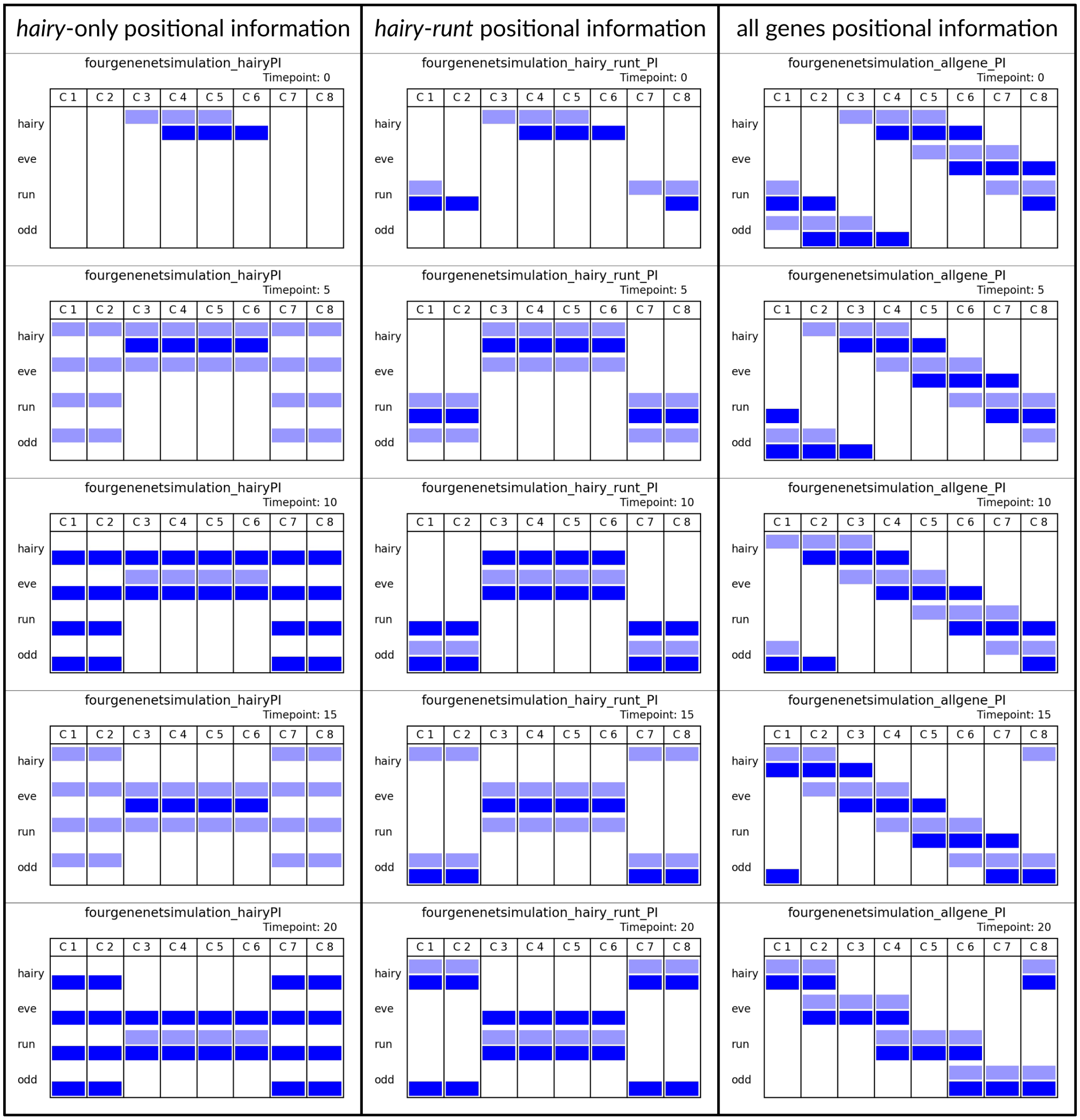


Figure S4: Four-gene network (Fig 4M) initialised with varying initial conditions. Only when expression of all genes is provided is a spatial, stable gene sequence formed.

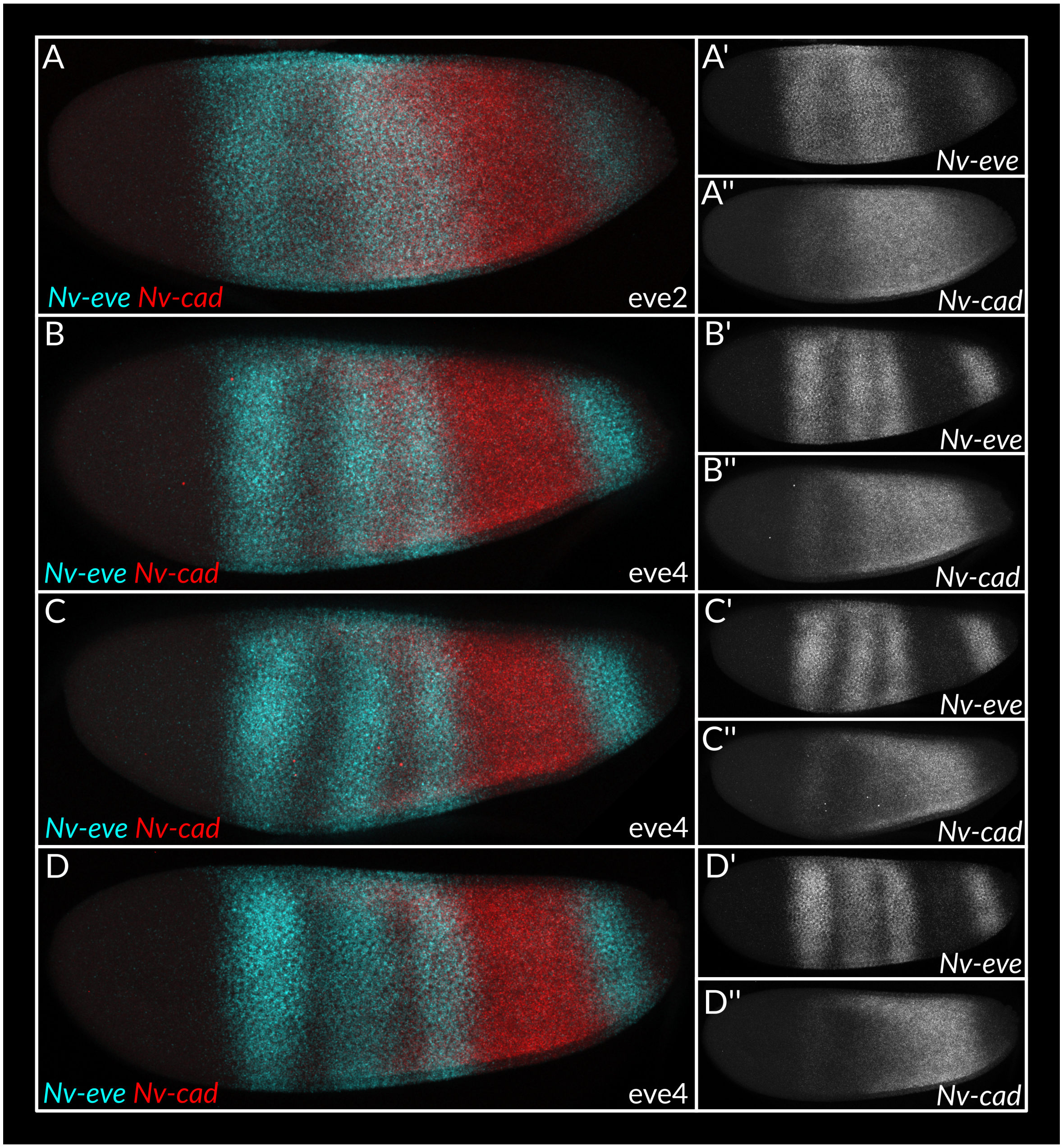


Figure S5: *Nv-cad* retraction during eve2 stripe splitting. All embryos are maximum intensity projections, anterior left and dorsal up, imaged under identical confocal settings, meaning that the strength of the *cad* gradient is directly comparable between embryos.

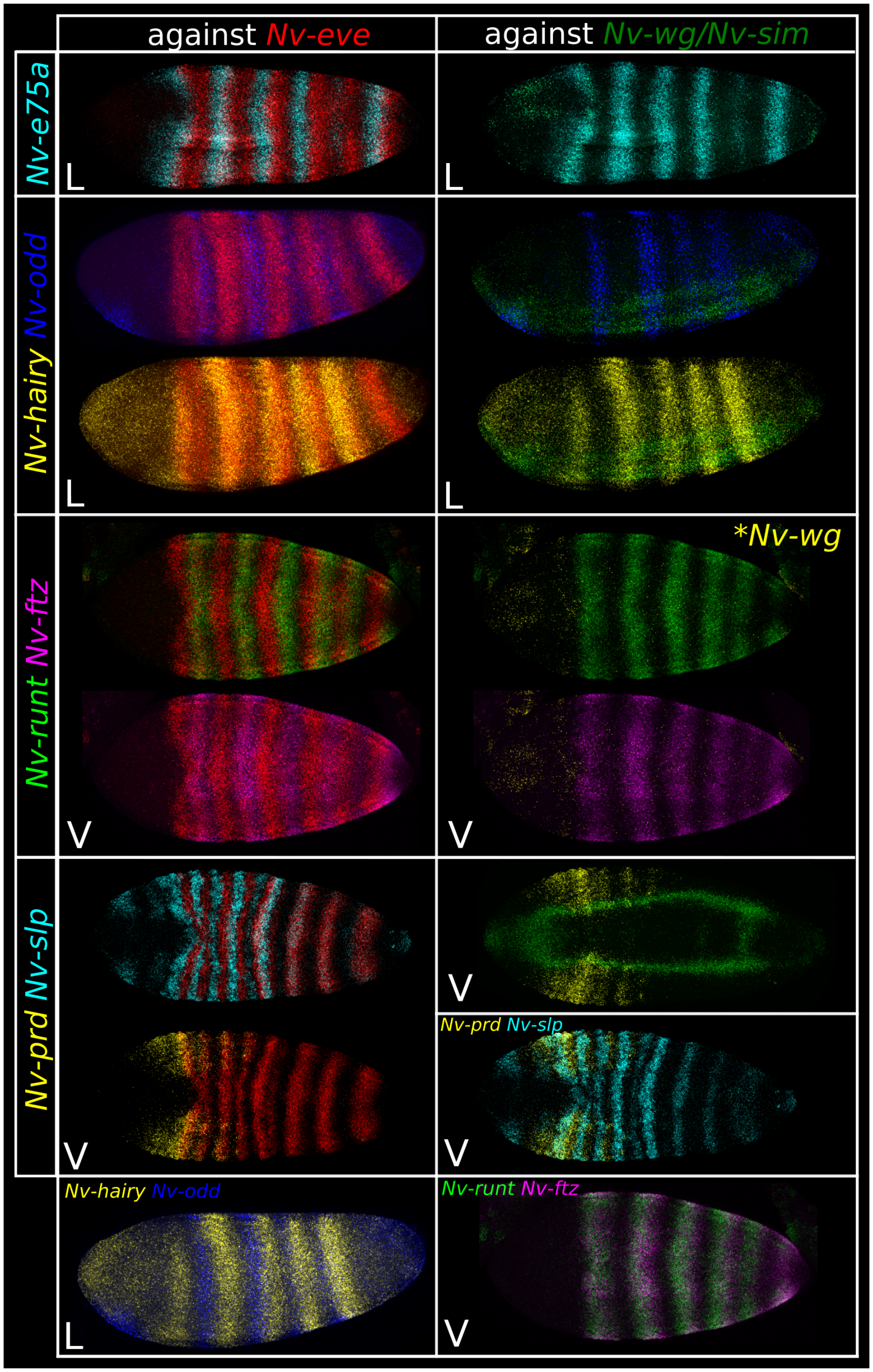


Figure S6: Expression of pair rule genes at the wg1 stage. All embryos are maximum intensity projections of half of the embryo (for clarity). Anterior left. Orientation is indicated by letters. L: lateral. V: ventral. D: dorsal.

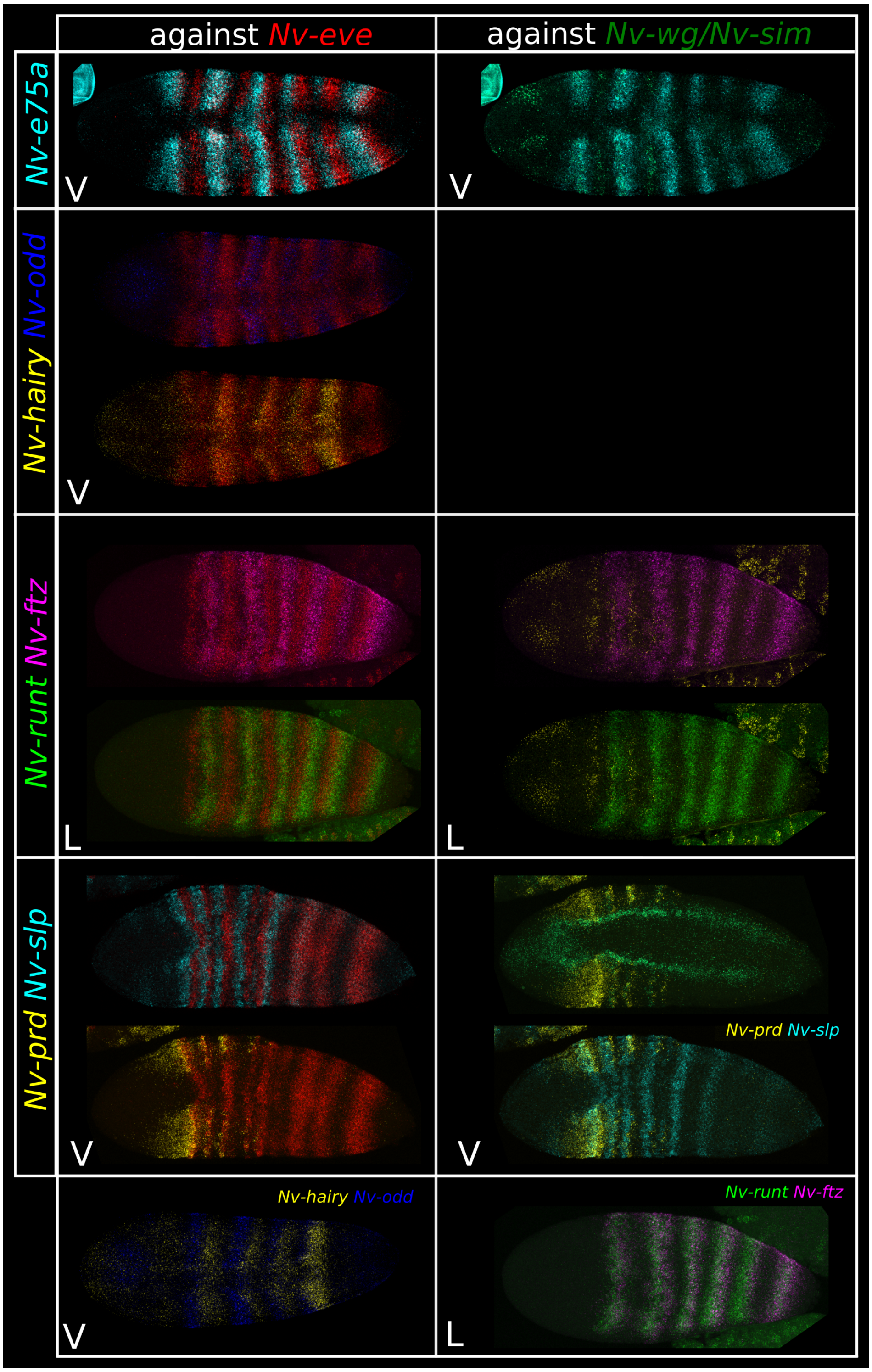


Figure S7: Expression of pair rule genes at the wg4 stage. All embryos are maximum intensity projections of half of the embryo (for clarity). Anterior left. Orientation is indicated by letters. L: lateral. V: ventral. D: dorsal.

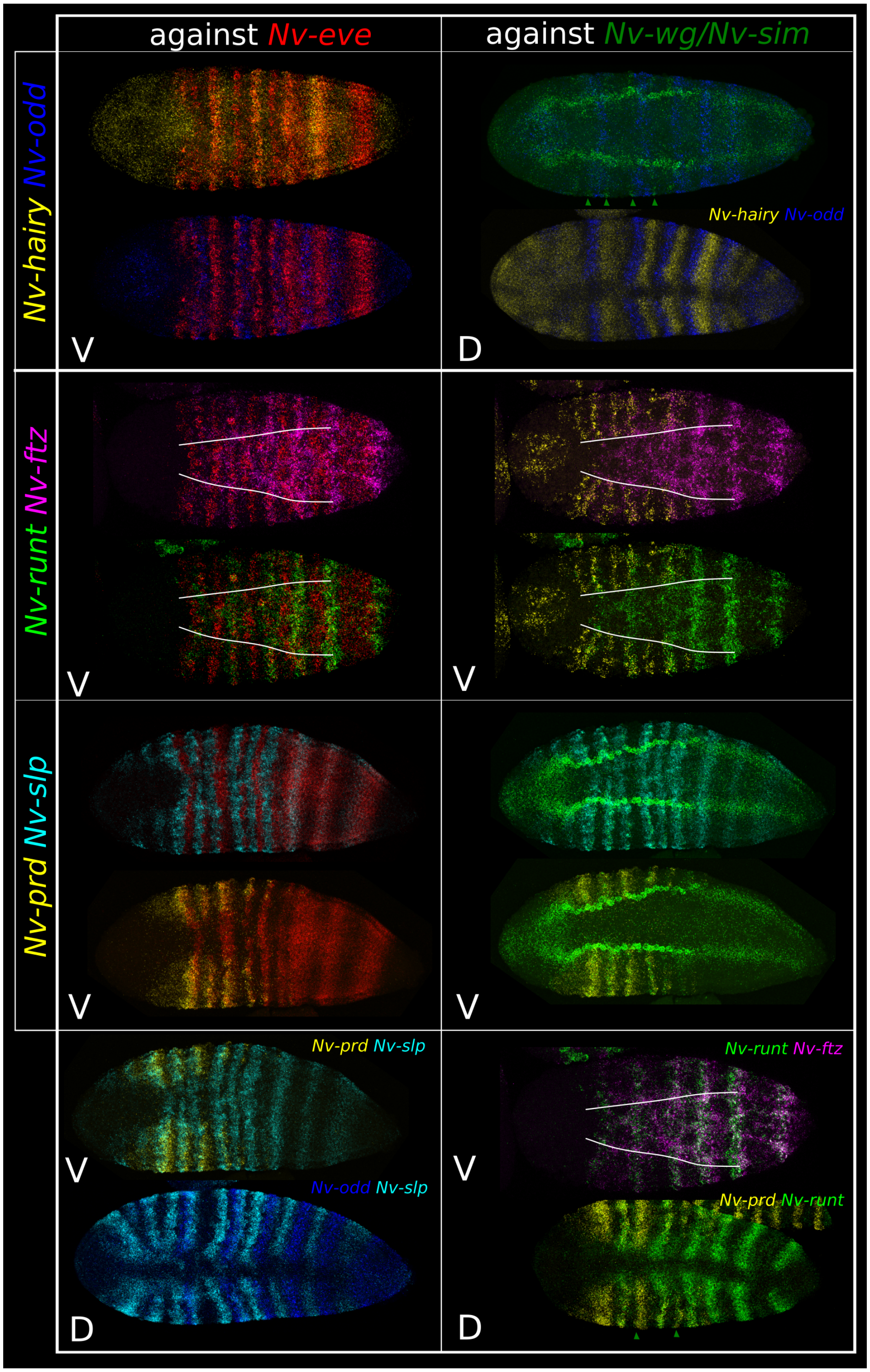


Figure S8: Expression of pair rule genes at the wg6 stage. All embryos are maximum intensity projections of half of the embryo (for clarity). Anterior left. Orientation is indicated by letters. L: lateral. V: ventral. D: dorsal.
